# A response to Yurko et al: H-MAGMA, inheriting a shaky statistical foundation, yields excess false positives

**DOI:** 10.1101/2020.09.25.310722

**Authors:** Christiaan de Leeuw, Nancy Y. A. Sey, Danielle Posthuma, Hyejung Won

**Affiliations:** Department of Complex Trait Genetics, Center for Neurogenomics and Cognitive Research, Amsterdam Neuroscience, Vrije Universiteit Amsterdam, Amsterdam, The Netherlands; Department of Child and Adolescent Psychiatry and Pediatric Psychology, Section Complex Trait Genetics, Amsterdam Neuroscience, Vrije Universiteit Medical Center, Amsterdam University Medical Center, Amsterdam, The Netherlands; UNC Neuroscience Center, University of North Carolina, Chapel Hill, NC, 27599, USA; Department of Genetics, University of North Carolina, Chapel Hill, NC, 27599, USA

## Abstract

Hi-C coupled multimarker analysis of genomic annotation (H-MAGMA) was initially developed to advance MAGMA by assigning non-coding SNPs to their cognate genes based on threedimensional chromatin architecture. Yurko and colleagues raised concerns that the SNP-wise mean gene-analysis model of MAGMA may allow inflation in type I errors. Accordingly, we updated MAGMA and found that the updated version (MAGMA v.1.08) effectively controls for error rate inflation. Intrigued by this result, H-MAGMA was also updated by implementing MAGMA v.1.08. As expected, H-MAGMA v.1.08 detected a smaller set of risk genes than its original version (v.1.07), but the overall statistical architecture remained largely unchanged between v.1.07 and v.1.08. H-MAGMA v.1.08 was then applied to genome-wide association studies (GWAS) of five psychiatric disorders, from which we recapitulated our previous findings that psychiatric disorder risk genes display neuronal and prenatal enrichment. Therefore, issues raised by Yurko and colleagues can be overcome by using (H-)MAGMA v.1.08.

---

We have recently introduced a computational framework, Hi-C coupled multimarker analysis of genomic annotation (henceforth H-MAGMA)^1^, that integrates functional genomic datasets (e.g. Hi-C) with a widely adopted GWAS annotation tool MAGMA^2^. The major concept of H-MAGMA is simple: it inherits the statistical framework of MAGMA in aggregating SNP-level association statistics into genes, while employing chromatin interaction profiles in linking SNPs to genes. We noted that this concept can be used in various settings, and mentioned that: *“H-MAGMA can be expanded into many different forms. For example, we decided to use MAGMA among many other tools available because it is most widely used; however, this framework is applicable to any other tools that convert SNP-level P values into gene-level association statistics…^1^”*

Yurko and colleagues have raised concerns about our work by pointing out that the statistical foundation of MAGMA exhibits error rate inflation when summary statistics are used as input for the SNP-wise mean gene analysis model^3^. Accordingly, MAGMA-based tools including H-MAGMA suffer the same issue. In response to this criticism, we updated MAGMA (MAGMA v.1.08, https://ctg.cncr.nl/software/magma) to control for the error rate inflation.

In MAGMA v.1.08, the SNP-wise Mean model was changed to use an alternative test statistic 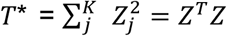, with *Zj* = Φ(*p_j_*) and Φ the cumulative normal distribution function; that is, *T\** is the sum over squared SNP Z-statistics. Jointly, for the vector *Z* we can assume *Z* ~ MVN(0, *S*), where *S* is the correlation matrix of the SNP genotypes. Given the eigendecomposition *S* = *QΛQ^T^*, we rewrite *Z* = *QΛ*^0.5^*D* for a random variable *D* ~ MVN(0, *I_k_*). It follows that 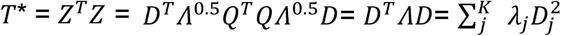, with *λ_j_* the *j*th eigenvalue and 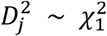. That is, the distribution of *T*∗ is equal to a mixture distribution of independent 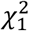 random variables.

Imhof^4^ provides a method of expressing the gene P-value *P* = *Pr(T\** ≥ *T_obs_\**) as an integral that can be evaluated numerically, which is the approach implemented for the updated SNP-wise Mean model (using Gauss-Kronrod quadrature for the integration). Whereas previously for *T* an approximate distribution was evaluated, in this Imhof approach for *T\** we directly evaluate the sampling distribution itself. Although this removes the problem that arose with the previous implementation, it is possible for the numerical integration to fail for some genes. We have therefore additionally implemented a fallback procedure that is used in case the numerical integration fails. This fallback procedure generates empirical P-values for those genes where the numerical integration fails, using an optimized simulation process to generate draws from 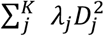. The number of simulations used is determined adaptively to reduce computing time, using up to a maximum of one billion simulations. Empirical P-values that are still zero afterwards are set to 0.5/1e9 = 5e-10. Though the actual P-value in such cases is likely to be significantly lower, running additional simulations beyond this becomes computationally infeasible. Any multiple-testing corrected significance threshold that is likely to be used will be generally much higher than this lower bound, and precision of the P-values around that threshold will be sufficient to reliably determine significance.

We performed a simulation study to evaluate the type 1 error rates of the updated model. This was based on the European cohort of the 1,000 Genomes data and the NCBI 37.3 gene definitions that can be found on the MAGMA site (https://ctg.cncr.nl/software/magma). The genes were first annotated to the data, resulting in a total of 19,248 annotated genes containing 9,220,220 unique SNPs, and 485.4 SNPs per gene on average. We duplicated individuals in the 1,000 Genomes data to attain a sample size of 5,000, and generated 1,000 null phenotypes drawn from a standard normal distribution, each of which was analyzed using the updated SNP-wise Mean model. For the output, we first estimated Bonferroni family-wise error rate FWER at *α* = 0.05 as the proportion of the 1,000 simulations for which at least one gene had a p-value below 0.05/19,248. This estimate came to 0.046, showing the FWER to be well-controlled. We then computed the type 1 error rate per individual gene at *α*’s of 5% and 0.1%. Quantiles of these per-gene type 1 error rates are shown in **Figure 1**, and as can be seen this distribution is consistent with the error rates being well-controlled.

**Figure 1.**
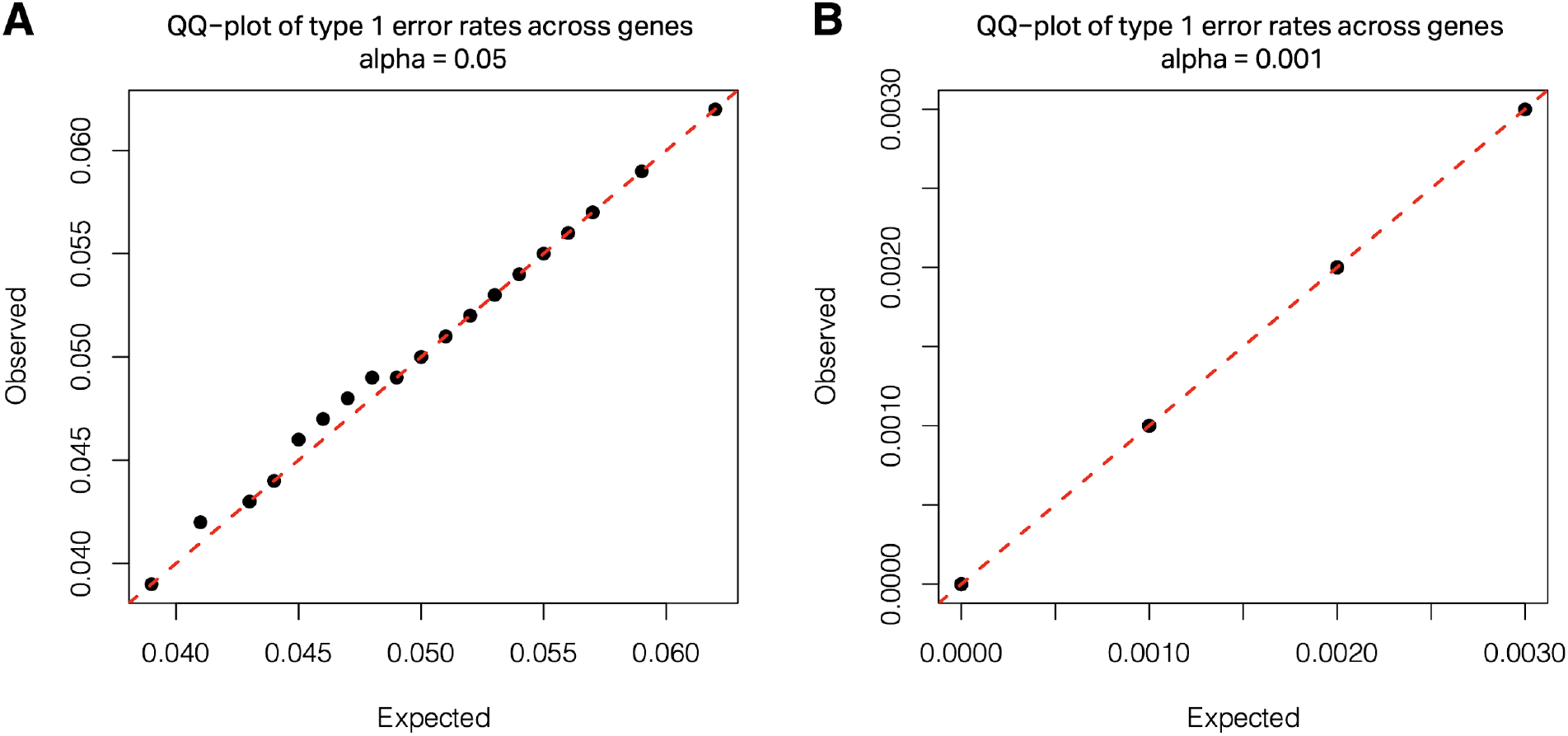
Distribution of per-gene type 1 error rates at significance threshold of 5% (**A**) and 0.1% (**B**). Quantiles (5th to 95th percentiles in steps of 5) are compared against the expected quantiles, obtained by taking the quantiles of a binomial distribution with 1,000 trials and a success probability of 5% or 0.1%, and dividing these by 1,000 (note that in **B** many of the quantiles overlap, and hence only four distinct points are visible).

Because Yurko et al., suggested that the error rate inflation is dependent on the gene size^3^, we also fitted a linear regression model (OLS), predicting per-gene error rate using gene size *S* (in number of SNPs per gene), density *D* (in estimated number of parameters per gene), and ratio *R* = *D/S*. In total, eight predictors were included: *S*, *D* and *R*, their squared values, as well as the log of *S* and *D*. This yielded an adjusted R2 of 0.000235 for the 5% error rates and of 0.0012 for the 0.1% error rates, indicating that there is no dependence of gene type 1 error rates on the size of, or level of LD in, a gene.

Finally, we re-analyzed 5 psychiatric disorder GWAS (attention deficit/hyperactivity disorder [ADHD], autism spectrum disorder [ASD], schizophrenia [SCZ], bipolar disorder [BD], major depressive disorder [MDD]) using H-MAGMA v.1.08 to replicate our initial findings in *Nature Neuroscience^1^*. MAGMA v.1.08 was used instead of MAGMA v.1.07 with the same code and input files as previously described^1^. Output files from running H-MAGMA v1.08 for all 5 psychiatric disorder GWAS are provided in the github repository (https://github.com/thewonlab/H-MAGMA).

Notably, the correlation between H-MAGMA v.1.07 and v.1.08 was over 0.99 (**Figure 2A**, Spearman correlation with Z-scores calculated from both versions), suggesting that the overall structure of association has not changed. As expected, H-MAGMA v.1.08 detected a smaller number of genes than H-MAGMA v.1.07(**Figure 2B**). Notably, the number of ASD risk genes identified by H-MAGMA v.1.08 (127) is similar to what Yurko et al. has reported (125)^3^, demonstrating that MAGMA v.1.08 can effectively control for a type 1 error rate. Since we used threshold-free, rank-based gene ontology (GO) analyses, we found that the GO results remain largely unchanged (**Supplementary Table 1**).

**Figure 2.**
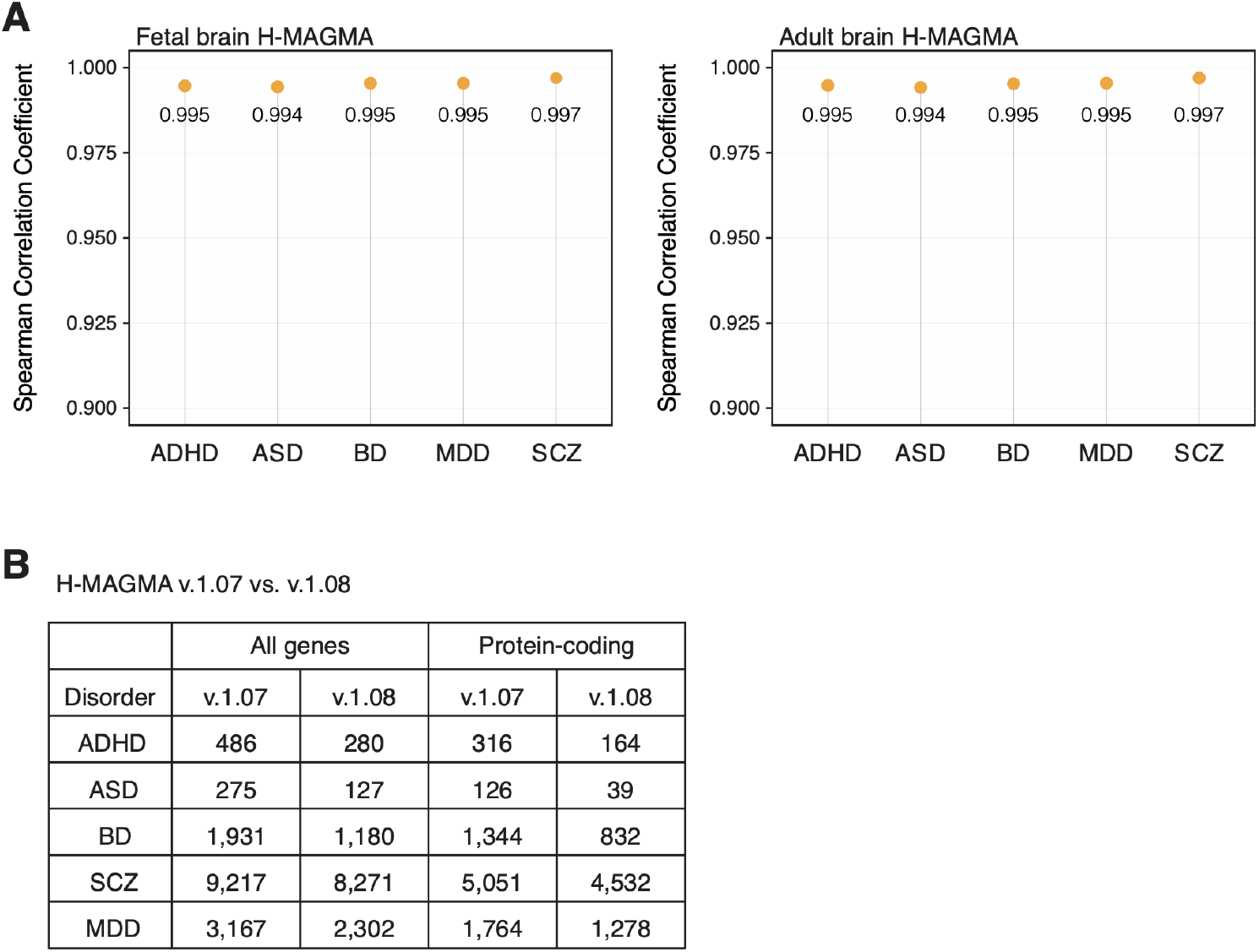
Comparison between H-MAGMA v.1.07 and H-MAGMA v.1.08. **A.** Spearman correlation coefficient between H-MAGMA v.1.07 and H-MAGMA v.1.08 for 5 psychiatric disorders in fetal brain H-MAGMA (left) and adult brain H-MAGMA (right). **B.** The number of psychiatric disorder risk genes (FDR<0.05) detected by H-MAGMA v.1.07 and v.1.08.

Moreover, we replicated our findings that psychiatric disorder-associated genes were highly expressed in excitatory neurons (**Figure 3A**). The only change we observed came from the developmental expression trajectories, because we no longer detected prenatal enrichment of ASD risk genes (**Figure 3B**). The rest of psychiatric disorders displayed prenatal enrichment, consistent with our previous findings^1^. Since ASD GWAS is less powered than other psychiatric GWAS, we believe prenatal enrichment of ASD risk genes will be recapitulated when a better powered GWAS becomes available. Indeed, ASD risk genes exhibited prenatal enrichment when we relaxed our threshold to FDR<0.2 (**Figure 3C**).

**Figure 3.**
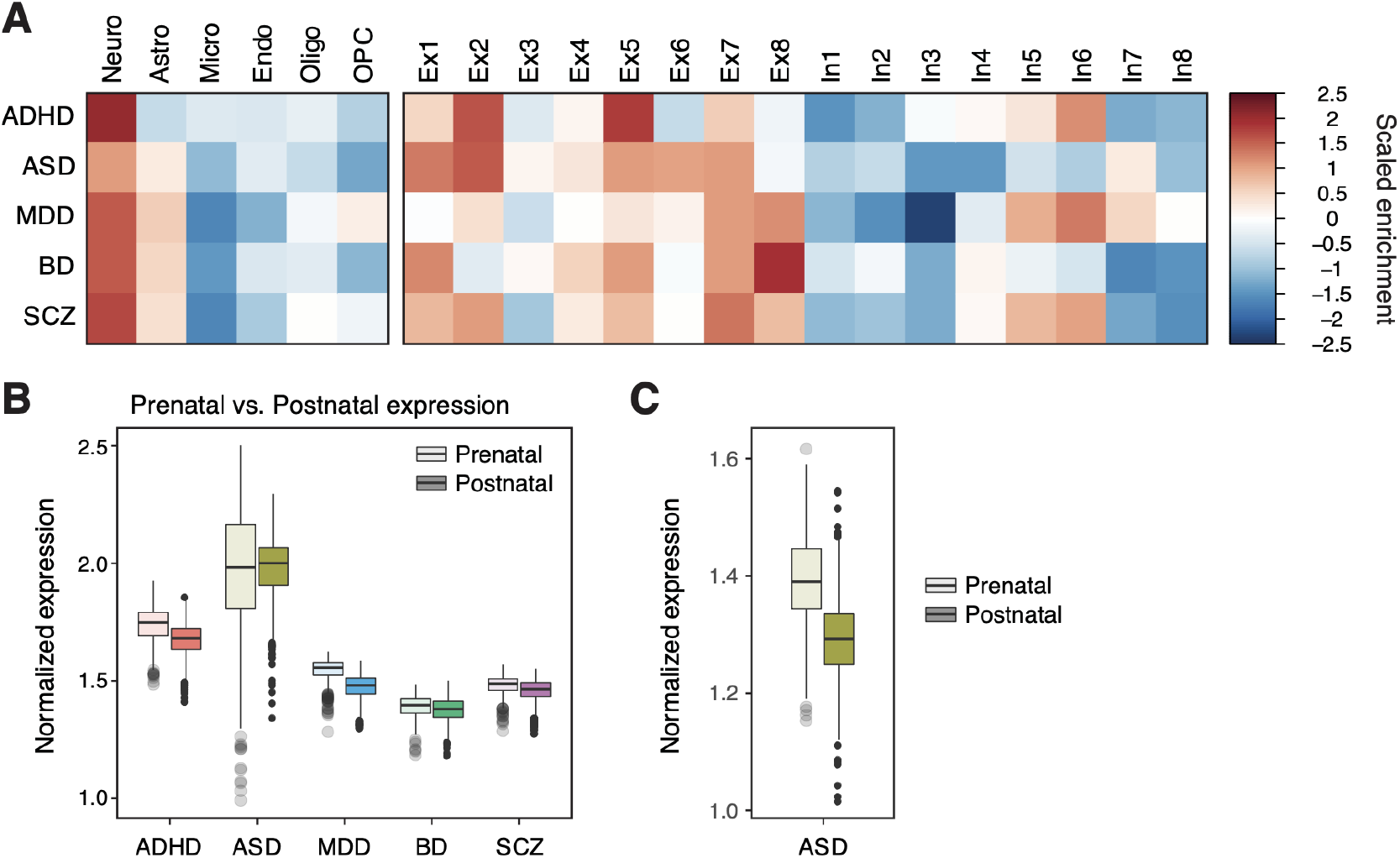
Cellular and developmental expression profiles of psychiatric disorder risk genes identified by H-MAGMA v.1.08. To define risk genes, FDR < 0.01 was used for GWAS with >20 GWS hits (SCZ, BD, and MDD), while FDR < 0.1 was used for GWAS with <20 GWS hits (ADHD and ASD). **A.** Psychiatric disorder risk genes are highly expressed in neurons than other brain cell types (left). Within neuronal subtypes, psychiatric disorder risk genes are enriched in excitatory neurons (Ex) than inhibitory neurons (In, right). **B.** Psychiatric disorder risk genes show higher expression values in prenatal brains than postnatal brains. The only exception was ASD. **C.** ASD risk genes defined based on FDR<0.2 show prenatal enrichment as other psychiatric disorder risk genes do.

In conclusion, the updated MAGMA (v.1.08) can effectively control for the potential inflation in the type 1 error rate. Tools that are built upon the statistical foundation of MAGMA, such as H-MAGMA, can overcome the recently reported issues^3^ by using MAGMA v.1.08.

## Notes

### Competing Interest Statement

The authors have declared no competing interest.

